# Ciliated cell domains with locally coordinated ciliary motion generate a mosaic of microflows in the brain’s lateral ventricles

**DOI:** 10.1101/2025.02.19.638730

**Authors:** Kennelia Mellanson, Lei Zhou, Tatiana Michurina, Anatoly Mikhailik, Helene Benveniste, Dimitris Samaras, Grigori Enikolopov, Natalia Peunova

## Abstract

Circulation of cerebrospinal fluid (CSF) through the brain’s ventricles is essential for maintaining brain homeostasis and supporting neurogenesis. CSF flow is supported by the structural polarization of multiciliated cells, which align with the flow direction. However, it remains unclear how the organization of tissue-wide polarity across the ciliary epithelium comprised of thousands of cells, determines the trajectory of the flow and efficient distribution of the CSF.

Here, we used new approaches to analyze the organization of translational polarity across extensive areas of the lateral ventricular wall. We also used live imaging to examine cilia motion, flow trajectories, and ciliary beat frequency (CBF) in live preparations of ventricles. In addition to the primary flow running across the ventricular wall from the posterior area to the anterior (P-A), we found multiple local microflows with both direct and curved trajectories that deviate from the mainstream P-A direction.

Our results suggest that the ciliated epithelium in the lateral ventricles varies in the alignment of ciliated cell translational polarity: whereas in the narrow dorsal area translational polarity is aligned with the direction of the mainstream flow, in the periphery of the mainstream it is organized into distinct cell clusters with locally aligned polarity vectors. We posit that the cluster organization of the multiciliated ependymal cells underpins the generation of a complex mosaic of flows, with the local microflows facilitating the wide spreading of the CSF across the ependyma. We demonstrate that nNOS is involved in the control of translational polarity, cluster organization, microflows, and CBF in the ependyma.

**Significance:** The flow of cerebrospinal fluid (CSF) is crucial for brain homeostasis. This flow is driven by the coordinated beating of cilia on thousands of ciliated cells. Planar polarity vector of the ciliated cells is aligned with the flow direction in the areas of the ependyma underlying the mainstream flow (1). However, the overall alignment of planar cell polarity and flow across the whole tissue is unclear. Here we report a discovery of complex flow patterns over the ependyma, consisting of numerous microflows in the periphery of the mainstream flow. Using new approaches, we found that planar polarity of the ependymal ciliated cells on the periphery of the mainstream flow aligns only locally, indicating a clustered organization of the ependyma that supports various flow directions.

## Introduction

Multiciliated cells, each bearing numerous motile cilia, are crucial components of the ependymal lining of the brain ventricles. The robust cooperative beating of cilia supports the flow of CSF throughout the ventricular system, thereby promoting the distribution of nutrients, growth factors, and metabolites throughout the brain, and participating in waste removal (2–6). CSF flow also contributes to the maintenance of the neurogenic niche on the lateral walls of the lateral ventricles and supports the migration of adult-born neuroblasts along the ventricular walls (7–11). Defects in ciliary function, whether inherited or acquired, can impede or disrupt CSF flow and waste clearance, and lead to a broad range of disorders; furthermore, these defects compromise neuroblast migration in the ventricular wall (6, 11).

The ependymal cell layer of the ventricular walls is a highly specialized tissue adapted for generating directional flow. Multiciliated cells exhibit several features of planar cell polarity (PCP) that define their structure and arrangement, as well as the overall architecture of the ependymal tissue. These cells display translational polarity manifested as clustering of ciliary basal bodies in patches on the apical cell surface, with the center of the patch shifted away from the center of the cell and toward the anterior side of the cell, in the general direction of flow (2, 12). The cilia also display rotational polarity, which defines the coordinated orientation of the basal bodies and axonemes of individual cells and of their neighbors in the direction of the ciliary beat. The collective activity of cilia across the lateral ventricles generates CSF streaming, directed from the posterior-dorsal part of the ventricle, where CSF is produced in the choroid plexus, towards the anterior horn, and then postero-ventrally toward the foramen of Monro where the flow enters the third ventricle. When examined in separate regions along the path of the flow, the median vector of the ciliated cells’ planar polarity in that region appears aligned with the direction of the CSF stream; this vector changes as the stream trajectory changes along the horn wall and the ventral aspects of the ventricle (8, 12)

Beyond supporting CSF flow, the arrangement of the multiciliated cells also shapes the neurogenic niche in the subventricular zone (SVZ) of the antero-dorsal area of the lateral walls of the lateral ventricles. Multiciliated cells surround B1 monociliated neural stem cells and form characteristic pinwheel-like rosettes (9, 11, 13–15). These stem cells give rise to progeny that gradually migrate toward the olfactory bulb and generate several subtypes of inhibitory neurons. The SVZ neurogenic niche is characterized by a unique planar vascular plexus, wherein dividing stem cells and their transit-amplifying progeny are found adjacent to the blood vessels of the plexus (16–18).

The basic PCP pathway is modulated by various intrinsic and extrinsic signals. Nitric oxide (NO), produced by various isoforms of NO synthase (NOS), is a versatile signaling molecule that affects multiple molecular targets and modulates broad range of physiological processes (19–21). The neuronal NOS isoform (nNOS) is expressed throughout the brain and participates in diverse signaling cascades, by activating guanylate cyclase or by directly modifying target proteins by nitrosylation (21, 22). Our previous studies have shown that an nNOS ortholog interacts with the components of the PCP pathway in the multiciliated cells of *Xenopus* embryonic skin (23). More recently, we have found that nNOS expressed in the multiciliated cells of the tracheal epithelium regulates the polarity of ciliated cells and the beat frequency of the motile cilia in the airway (24).

In this study, we explored the link between the tissue-level translational polarity of the multiciliated cells in the ependyma, the direction of cilia motion, and the flow direction across the ventricular wall.

In our experiments with the fixed tissue, besides the areas in which translational polarity vectors are aligned with the dominant posteroanterior (P-A) direction of the flow, we detected areas where cells are aligned by their translational polarity vectors only locally with their immediate neighbors, forming clusters with different mean vectors of translational polarity in each cluster. In our live imaging experiments, when examining extended regions of the ventricular explants, we found that besides the mainstream flow with a straight trajectory, generally running in P-A direction, there are multiple local flows with the trajectories sharply different from the main stream. We posit that the diversity of cell clusters with locally aligned polarity vectors may underlie the observed mosaic of local flows, contributing to the efficient distribution of CSF across the ventricular wall.

We also focused on the role of nNOS in the function of the ciliated cells, generation of the flow, and in neurogenesis. Our results indicate that nNOS is required for several key features of ependymal cells: the establishment of ciliated cell planar polarity, organization of cell clusters with locally coordinated ciliary motion, and the maintenance of CBF, thereby supporting robust directional flow along the ventricular walls. Furthermore, we show that nNOS regulates the progenitor proliferation pattern in the neurogenic niche of the SVZ.

## Results

### nNOS controls the cell size and shape of the multiciliated ependymal cells of the brain ventricles

We investigated the presence and distribution of the nNOS isoform in the ventricular ependyma of the adult mouse brain by immunohistochemistry. nNOS signal was detected primarily in axonemes of the ependyma (Fig. 1A, B, C). This observation was supported by the analysis of the nNOS-CreER/Ai9 mouse line (25): after inducing nNOS-driven recombination with tamoxifen, we observed a dTomato signal in the ependymal cells of the lateral ventricular wall. Co-staining with antibodies to dTomato, acetylated alpha-tubulin (a ciliary marker) and phalloidin (to highlight the apical actin cytoskeleton) revealed dTomato signal on the apical cell surface of the ciliated cells, concentrated in the tufts of cilia, implying nNOS expression in the multiciliated cells of the ventricles (Supplementary Fig. 1).

**Figure 1.**
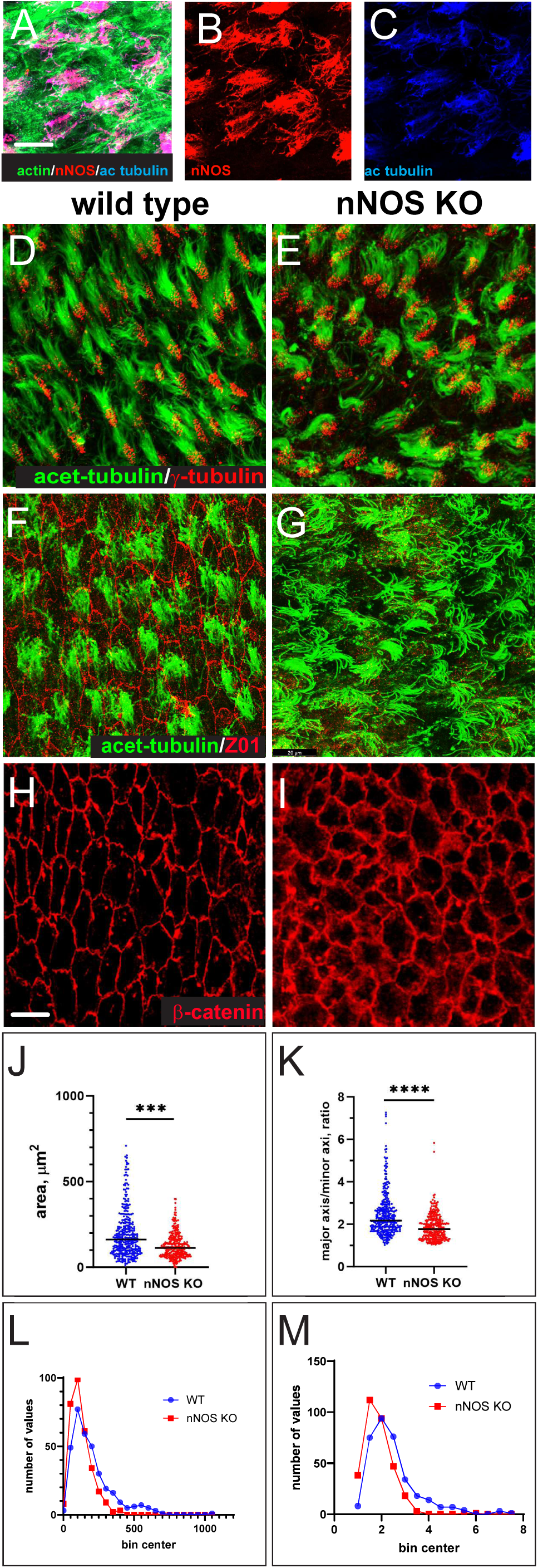
The structure of the multiciliated brain ependymal cells is affected by the deletion of nNOS gene. A-C Expression of nNOS in the ciliated cells. Immunochemistry with antibodies to nNOS (red), acetylated tubulin (blue) and staining with phalloidin to reveal actin in apical cortex(green) antibody indicated the expression of nNOS in the ciliated cells. Scale bar is 10 µm for (A-C). **D, E** There is no changes in the length or density of cilia in the multiciliated cells in the brain ependyma of nNOS-null (nNOS KO) mice.. The explants are stained with antibodies to acetylated α-tubulin (green), and γ-tubulin (red). **F-I** Changes in shape and size of multiciliated cells apical surface in the brain ependyma of nNOS KO mice. **F-I** Apical surface of the wild-type (F,H) and nNOS KO (G,I) ependymal cells, labeled by antibodies to ZO1 (red)and acetylated tubulin ( green) and B-catenin (red) Scale bar is 20 µm. **J, L** The area of the apical surface is smaller in the nNOS KO than in the wild-type (data presented as average (J) and as frequency distribution (L) (blue dots and blue line in wild type and red dots and line in nNOS KO ependyma). **K, M** Apical surface of the ciliated cells of wild-type ependyma have elongated shape (H,K). Ciliated cells in the ependyma of nNOS KO are rounder (I,K). The change of elongated shape in the ciliated cells apical surface in nNOS KO ependyma is quantified as ratio of the long to the short cell axes, presented as average (K) and as frequency distribution (M) in wild-type and nNOS KO ependyma.

We then examined the polarity of the ciliary orientation, the morphology and polarity of the ciliated cells in the ependyma, and the ventricular tissue architecture, sampling various areas in the ventricles of the wild-type and nNOS knockout (nNOS KO) animals. Freshly excised brain ventricles were flattened, fixed, and stained with molecular markers of the cilia (acetylated β-tubulin), the basal bodies (ODF2 or γ-tubulin), the apical surface (β-catenin), or marker of tight junctions (Z01). The processed whole-mount ventricle samples were oriented stereotypically horizontally on slides to maintain the reference of the planar cell polarity features relative to the A-P axis of the brain. Several areas of the ventricles were routinely photographed, and data for the groups of the wild-type and nNOS KO littermates were collected for statistical analysis (each group contained at least 5 animals).

Cilia axonemes in the ependyma of the wild-type and nNOS KO mouse littermates did not differ in length (Fig. 1D, E) and ultrastructure, as revealed by electron microscopy (not shown).

However, the changes produced by the loss of nNOS were evident in the overall shape and arrangement of the ependymal cells (Fig. 1F, G). Antibodies to β-catenin and Z01 revealed the characteristic elongated shape of the ciliated cells in wild-type ependyma, with consistent long and short axis ratios (Fig. 1H, K). Both the shape and size of the ciliated cells and their arrangement within ependyma were altered in the ependyma of nNOS KO animals. The apical surfaces of ciliated cells in the mutants were smaller (Fig. 1H, I, J) and rounder, with the ratio of the long to short axes of the apical cell surface lower than observed in the wild type (Fig. 1K). These findings were supported by the frequency distribution analysis of wild-type and nNOS KO ciliated ependymal cells, which suggested that the observed changes were consistent in the majority of analyzed cells (Fig. 1L, M).

### Arrangement and orientation of multiciliated cells in the ependyma and the role of nNOS

In the wild-type ependyma, neighboring multiciliated cells appeared aligned along their long axes, oriented orthogonally relative to the A-P body axis. (Fig. 1H, 2A). This characteristic feature was notably altered in the ependyma of nNOS KO mice, with noted disruption of the alignment and arrangement of the ciliated cells in the tissue (Fig. 1I, 2B).

**Figure 2.**
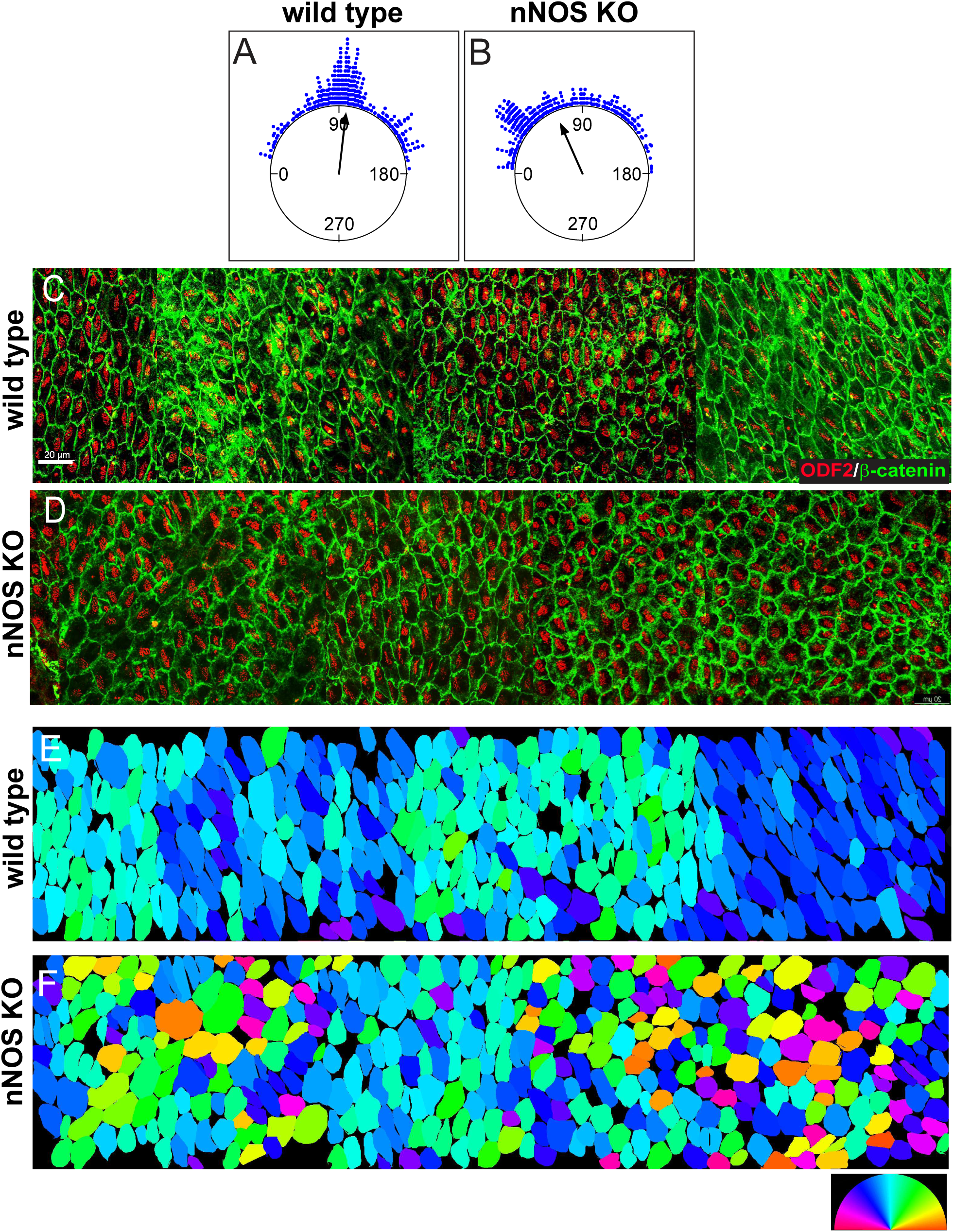
Changes in the orientation multiciliated cells affects the architecture of ependyma of nNOS KO animals. **A-D** Ciliated cells in the wild-type ependyma show alignment of orientation of their apical surface long axis relative to the A-P axis of the brain (A,C). The direction of the arrow represents the mean vector for the entire field, and points to 96°, almost orthogonally to the A-P axis. The arrangement of the ciliated cells in the nNOS KO in the ependyma is altered (D)and their apical surface’s orientation relatively the A-P axis is shifted to 69°B). Length of Mean Vector (r) in the wild-type - 0.865; (r) in nNOS KO-0.771. n=198 cells analyzed for the wild-type and 217 cells for the nNOS KO ependyma, number of animals n=5 for each genotype. Staining: β-catenin (green), ODF2 (red). C-F. Assessment of the long axis alignment across the ependyma in the heat map style diagram. **C, D** Extended area of ependyma (stitched images of neighboring fields) of the multiciliated cells in the wild-type (**C**) and nNOS KO (**D**) ependyma. Staining ODF2 (red) and the b-catenin (green). Scale bar is 20 µm. **E, F** Heatmap rendering of the ciliated cells apical surfaces orientation relatively A-P axis in color: the wild-type field (E) and in the nNOS KO field (F). Color ring scale is 0-180°.

We next sought to visualize *in situ* the alignment and orientation of the ciliated cells’ apical surfaces relative to the A-P axis in the extended area of ependyma. We stitched the images of four or five neighboring areas into composite panels (Figure 2C, D). We developed an algorithm to find the long axis of each ciliated cell in the image and to calculate the angle between the long axis of the elongated apical surface of each ciliated cell and a reference line across the examined area, (which coincided with the A-P axis). After the angle for each cell in the examined area was calculated, the angle values were represented using a standard color ring scale. This approach yielded heat map-style diagrams of the angle value distribution (Fig. 2E, F), which preserved spatial information and provided color-coded numerical information for each cell orientation across large tissue areas.

The overall distribution of angles was relatively narrow in the wild type ependyma (Fig. 2 E). In contrast to the relatively uniform arrangement and orientation of cells lining the ventricular wall in wild type ependyma, the cell arrangement in nNOS KO mice varied significantly and contained large areas with poor or absent axis alignment and orientation (Fig. 2F). Remarkably, despite the pronounced orientation changes in the mutants, within preparations from the same ventricles, we observed interspersed regions with a “normal” phenotype, comprising an axis orientation distribution resembling that of the wild-type.

These results uncovered new features of the tissue-level polarity in multiciliated ependymal cells within the lateral ventricles, emphasizing a consistent alignment of the long axes of ciliated cells’ apical surfaces. These axes are orthogonal to the anterior-posterior (A-P) axis of the brain. Notably, the deletion of nNOS led to changes in the size and shape of their apical surfaces (Fig. 1J, K) and misalignment relative to the A-P body axis in the ependyma (Fig. 2E, F).

### Translational polarity of the multiciliated epithelium and the role of nNOS

In the ependymal ciliated cells, basal bodies aggregate into patches that cover only a portion of the cell surface and are shifted toward the cell’s anterior side (12, 26). The shift is characterized by the vector from the center of the cell toward the center of the patch (“patch vector”) and by the distance between the center of the patch and the center of the cell (“patch displacement”) (26). Tissue-level translational polarity is manifested as consistent alignment of the patch vectors among adjacent cells.

The feature of translational polarity was evident in groups of neighboring cells (Fig. 3A, B). The data, collected across the ependyma, presented as circular plots, showed a distribution of the translational polarity vectors in the wild-type mice, with a standard deviation (SD) of 57°. The distribution was significantly broader in nNOS KO, with an SD of 71° (Fig. 3C,D). We computed the patch vector and patch displacement (Fig. 3C-H) for each cell, as previously described (Boutin et al., 2014). The ratio of the patch to cell areas was higher in the mutants, possibly reflecting a smaller apical area of ciliated cells (Fig. 3E, G). The patch displacement values were significantly lower in the mutants than the wild-type (Fig. 3F, H).

**Figure 3.**
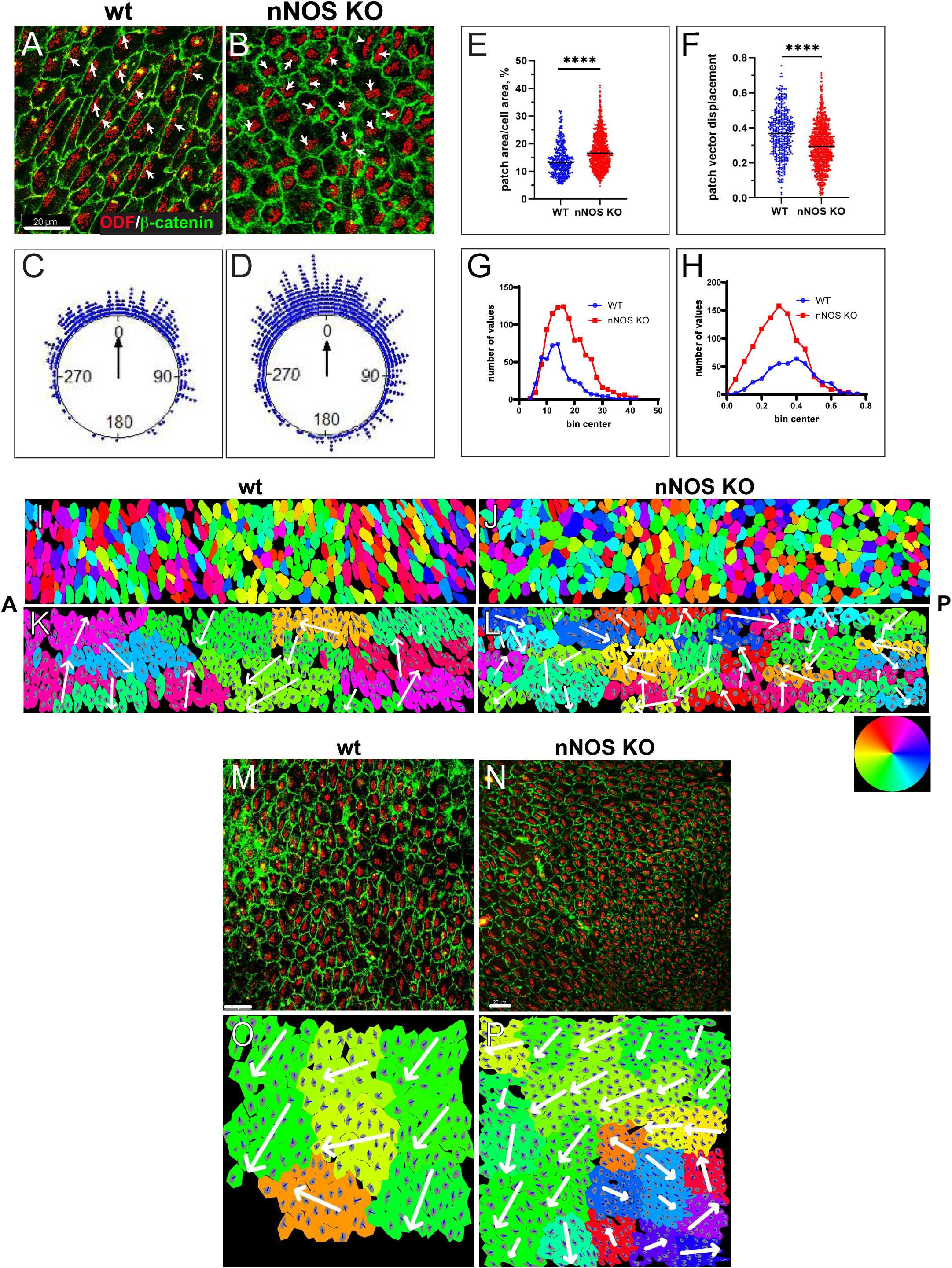
The translational polarity of the multiciliated cells in the wild-type and nNOS KO ependyma. **A B** The samples of the wild-type (A) and nNOS KO (B) ependyma with the cell walls and basal body patches labeled with antibodies to β-catenin (green) and ODF2 (red), respectively. Scale bar is 20 µm. Arrows indicate the vectors of translational polarity. **C, D** The distribution of the patch vectors in wild-type (C) and nNOS KO(D) ependyma. The dots at the perimeter of the plot represent the angle between the patch vector in the single cell and the average patch vector in the area. n=423 cells, 25 areas were examined for the wild-type and 975 cells from 36 areas for the nNOS KO ependyma: n=5 number of wild-type and n-5 of nNOS KO genotype. **E, G** The patch area occupies a larger fraction of the apical surface in the nNOS KO ciliated cells than in the wild-type. **F, H** Patch displacement vectors are shorter in the nNOS KO ciliated cells than in the wild-type. Data are presented as a scatter plot **(E, F**) and as frequency distribution **(G,H**) for the wild-type (blue dots and line) and nNOS KO (red dots and line). **I, J** Heatmap diagram of the distribution of the translational polarity vector in the wild type (I) and nNOS KO (**J**) ependyma. The translational polarity vector in each cell is expressed in color as angle between the patch vector and the reference line (A-P brain axis), color is based on the corresponding position of the angle in the standard color ring (original images of the wild-type and nNOS KO are in Fig. 2C and 2D, respectively). **K, L** Neighboring cells, that showed alignment of their translational polarity vectors are depicted as clusters. Each cluster is characterized by the alignment of translational polarity vector, with SD of 60° (the value is chosen as close to the observed value for the wild-type; Fig. 3G). The entire cluster is labeled by the color corresponding to the value of the mean vector, based on the standard color ring (See: Methods /Algorithm cell clusters /alignment by translation polarity vector). There are 12 clusters in the wild-type and 27 in the nNOS KO. **K** In the wild type besides cell clusters with translational polarity aligned with the dorsal posterior direction of the flow (yellow, green clusters) there are clusters of cells aligned along the vector directed elsewhere, poised to generate flow in alternative directions. In nNOS KO animals’ sample of same size ependyma contains larger number of clusters 27 aligned along larger variety of directions and clusters contain fewer cells than in the wild-type, where there are only 12 cell clusters (K,L).

To visualize the tissue-wide translational polarity *in situ*, we examined the translational polarity vector distribution in the composite panels of images (the same as used for analysis presented in Fig. 2C, D). Specifically, we computed the patch vector for each cell, then computed the angle between the patch vector and the reference line (the A-P axis), expressed the angle’s value on the standard color scale, and assigned a corresponding color to each cell in the composite panel. This process produced a heatmap diagram of the translational polarity vector distribution across the entire analyzed tissue areas (Fig. 3I, J).

Realizing the statistical nature of the alignment of translational polarity vectors in the ependyma, we next sought to characterize the vector alignment between neighbors *in situ* in the sampled area. We developed an algorithm that identifies neighboring cells whose patch vectors align within a variable value of standard deviation (SD). This approach revealed that *in situ* distribution of translational polarity vectors in the ependymal tissue was significantly more complex than indicated by the circular plots. For instance, setting the algorithm to identify neighboring cells whose vectors align within a 60° SD revealed several clusters in the wild-type ciliated tissue. Each cluster consisted of cells aligned to a mean vector, with a standard deviation of 60° within the cluster. However, cells in adjacent clusters were aligned along vectors in different directions, as shown in Figure 3K. This indicates that in the sampled area of the wild-type ependymal tissue cells are aligned by translational polarity vectors only locally, within clusters of neighboring cells. The largest cluster in the wild-type panel consisted of cells aligned in the general P-A direction of the flow (yellow-green parts of the color ring); however, numerous other clusters, including those immediately adjacent to the large cluster had significantly different median translational polarity vectors (Fig. 3I, K). Such cell cluster arrangement suggests that the flow might be changing its direction when passing through these tissue domains. Summarily, such complex pattern of the tissue-wide translational polarity may lead to the flow of a more complex trajectory than the general P-A direction of the dorsal stream.

The loss of nNOS resulted in the reorganization of the cell clustering such that the cells aligned along their patch vector formed smaller clusters, thereby increasing the number of clusters (Fig. 3J, L). This suggests that in the nNOS KO, the resultant flow over the area of the same size as in the wild-type might change its direction more frequently than in the wild-type because of a larger number of clusters, each with a different vector of translational polarity. (Fig. 3J, L).

The alignment of the translational polarity vectors tested in the ventricular areas underlying the mainstream flow was highly consistent, with an angular deviation (SD) of less than 45° (Fig. 3L.M). A distinct feature of the ependyma underlying the mainstream flow in nNOS KO mutants was the presence of numerous cell clusters with locally aligned vectors of translational polarity forming swirl-like structures, thus disrupting the global translational polarity alignment (Fig. 3N, O). This finding suggests that even in the areas underlying the mainstream flow the deletion of nNOS led to the misalignment of translation polarity with the flow direction.

### Roles of nNOS in flow in lateral ventricles

In the lateral ventricles, CSF, produced by the choroid plexus in the posterior part of the ventricle, flows anteriorly through the dorsal corridor, bends ventrally after reaching the anterior horn, then moves ventro-posteriorly toward the foramen of Monro, and enters the third ventricle (8). We investigated the relationship between ciliary motion and flow trajectories through live imaging of both ciliary motion and flow velocity in the explants, by tracking the movement of fluorescent beads. We stained the dissected ventricles with Fast Red 640, which allowed us to visualize the cilia and to follow the direction of ciliary motion. We also added Alexa488-linked 1µm microspheres in the viewing chamber, which enabled tracking of bead movement in the recordings, and analysis of the flow velocity and trajectories. Moreover, we determined the CBF distribution *in situ* over the same areas by using custom-made software, which yielded heatmaps indicating CBF variation across the field on a 0–100% normalized scale. Recordings started at a consistent site above the choroid plexus, where fluid flow was easily visualized, then proceeded toward more anteriorly and laterally positioned areas, sampling the direction of the ciliary motion, flow direction and trajectory, and CBF.

In the posterior areas of the dorsal corridor, the CSF flow showed a straight, anteriorly oriented trajectory (Fig. 4A-C and Supplemental movie S1), as reflected in the movements of the beads across the dorsal corridor in P-A direction, and the cilia demonstrated coordinated motion, albeit with varying CBF across the field (Fig. 4B, C). Inspection of the more anteriorly and laterally positioned areas revealed more complex flow patterns, with local microflows that changed directions abruptly; made numerous detours dorsally and ventrally, away from the general stream of posterior-anterior direction; and showed both straight and curved trajectories (Fig. 4D-E, G-H, S1-S4).

**Figure 4.**
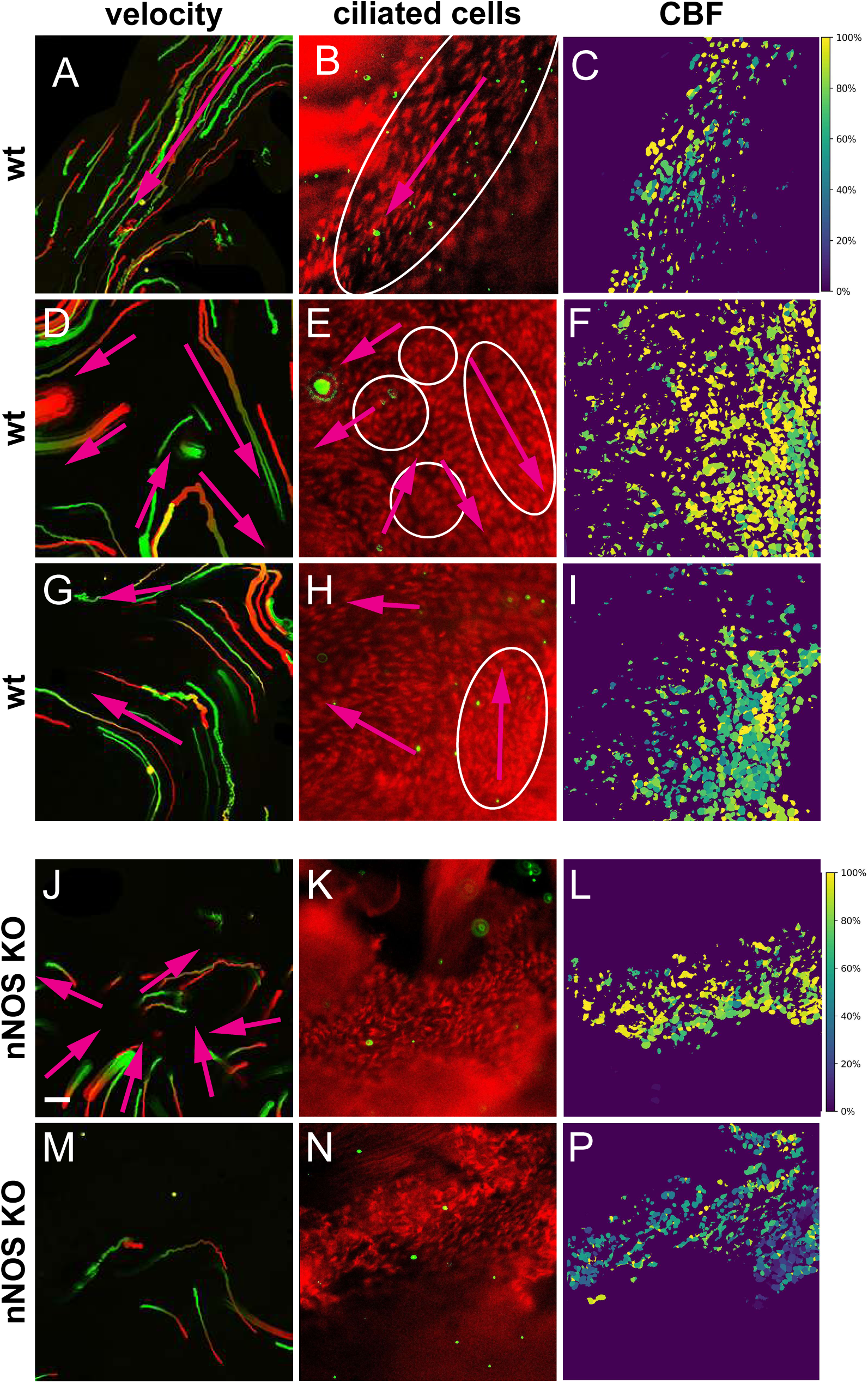
Flow and CBF in the wild-type and nNOS KO ependyma. Flow directionality (left) as represented by the beads’ tracks; the frame of the movie, captured in live imaging, middle panel); and the distribution of CBF (right panel) for the same field (each row). Three areas of the wild-type ependyma with straight and curved local flow (A-C, D-I)) and corresponding movies S1-S3, S4), and two areas of the nNOS KO ependyma with variety of local flows (J-L, M-P) and corresponding movies S5-S6,S7) are presented. The color of the tracks reflects the direction of the beads’ movement (red is the front end). Regions of local flows with distinct directionality, the ciliary beat direction in the wild type are marked by ovals and arrows. Scale bar is 20 µm for (A-P).

Analysis of the ciliary motion indicated that ciliated cells with coordinated ciliary motion were organized into domains, each with distinct ciliary beat orientation, this feature presumably underlying the local microflows with distinct bead trajectories (Fig. 4D-E, G-H, S1-S4).

Neighboring domains often produced local microflows that differed in direction (Fig. 4E, H, S1-S4). Of note, we did not detect differences in CBF between regions with straight or convoluted microflows (Fig. 4C, F, I).

In the nNOS KO ventricles, we observed several deviations from the wild-type patterns (Fig. 4J, M and Supplemental movies S5-7). The distinct nNOS KO-specific features included areas with turbulent flow, marked by short chaotically directed bead trajectories oriented towards the same point, suggesting a collision of the microflows (Fig. 4 J). Notably, although the cells within these areas showed normal CBF, their cilia lacked beat coordination with those of neighboring cells, thus resulting in cilia beating against each other (Fig. 4K, Supplemental movies S5, S7). Furthermore, we found isolated areas with substantially slowed CBF and no ciliary beat synchrony, even within individual cells in these areas (Fig. 4M-P and Supplemental movies S8, S9).

Together, our results suggest that besides the general fluid stream running in the P-A direction, the overall flow through the dorsal corridor comprises numerous local microflows exhibiting a mix of both straight and curved patterns. We posit that the complexity of these flow patterns is underlaid by the domains of ciliated cells in which ciliary movement is coordinated locally within the domain but not between different domains. Our results also suggest that loss of nNOS/NO affects planar cell polarity of the ciliated cells, CBF, domains’ composition in the ependyma, and flow directionality across the ventricular walls. Thus, diverse aspects of nNOS function contribute to both structural organization and planar polarity of the ependymal tissue and to the dynamics of cilia motion, coordination of the motion, and flow velocity and directionality.

### Effects of nNOS on the neurogenesis in the lateral ventricles

The lateral walls of the lateral ventricles are critical sites of continuous cell proliferation throughout the animal’s lifespan. B1 cells, located at the centers of multiple pinwheel structures, act as neural stem cells and generate neuroblast progeny that migrate toward the olfactory bulb and ultimately differentiate into olfactory neurons. Stem cells within this niche possess basal feet that contact blood vessel endothelial cells and potentially facilitate signal exchange between the vasculature and stem cells, thereby contributing to the regulation of neurogenesis (11, 13). The vasculature in these neurogenic regions exhibits a distinctive pattern, forming a plexus of vessels running closely alongside the ventricular walls. Importantly, the emergence of new neuroblasts occurs close to blood vessels (16), (18).

The ventricular niche vasculature which regulates NSC proliferative activity can be affected by various factors, including mutations and aging (27). Therefore, we compared the ventricular wall blood vessel arrangement between wild-type and nNOS KO mice by using an intravital dye, detecting no apparent differences (Fig. 5A, B). We then analyzed blood vessels tortuosity, assessed as the ratio between the measured length of the vessels’ path with its twists and turns and the length of the straight line of the most direct path (26), which similarly did not show apparent differences (Fig. 5E, F)

**Figure 5.**
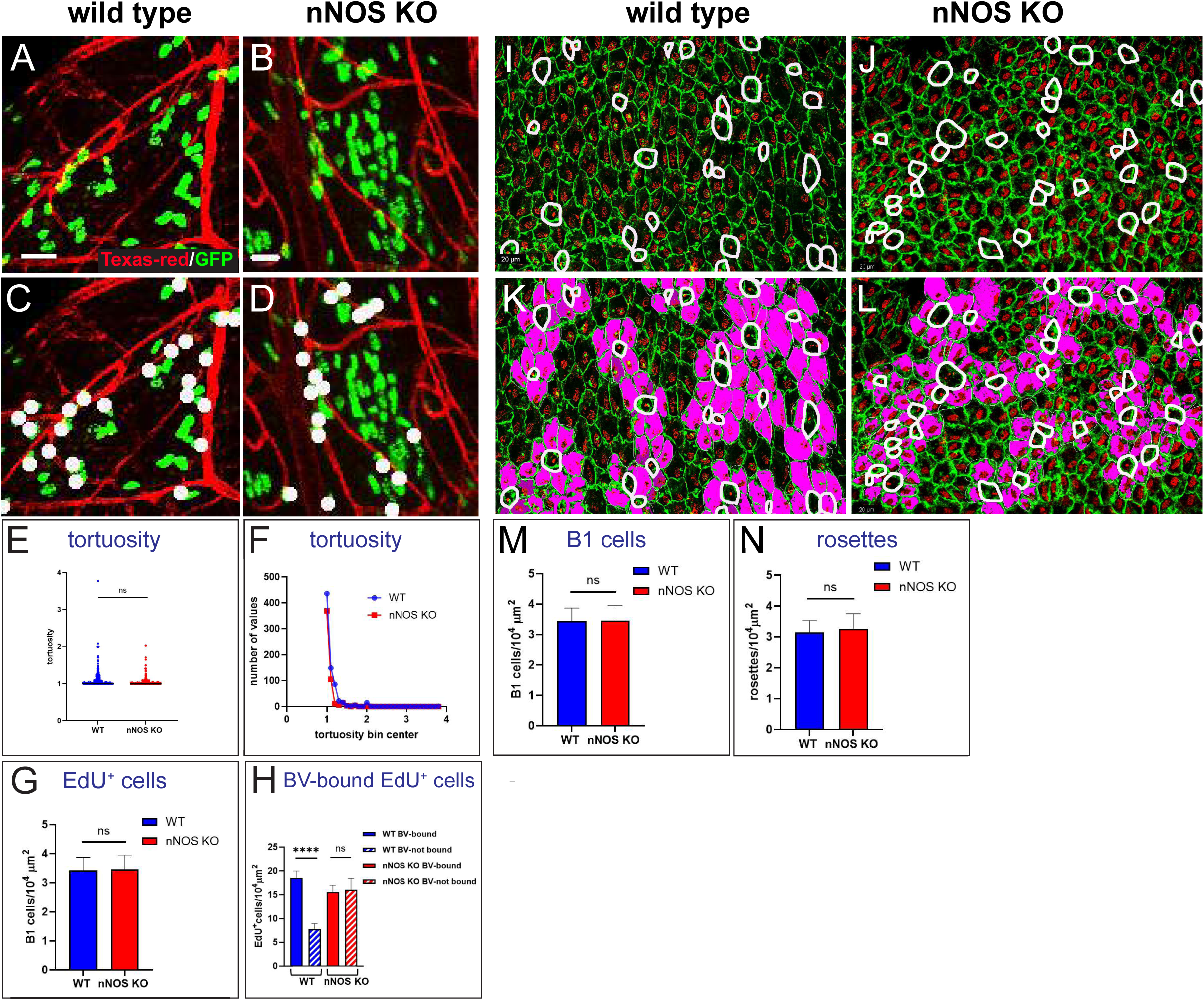
Proliferation of neural progenitor in relation to the blood vessels in the wild-type and nNOS KO ependyma. **A, B** Distribution of dividing cells in relation to the blood vessels in the SVZ. The characteristic plexus of the blood vessels is visualized with Texas Red lectin (red) and dividing cells are marked by the incorporation of EdU (green). Note the close apposition of the labeled nuclei of the dividing cells to the blood vessels in (A, B)labeled by white dot for cell counting (C,D) and numerous dividing cells not in contact with the blood vessels in left green for cell counting ( C,D). Scale bar is 20 µm. **E,F** Arrangement of the blood vessel branches, as reflected in their tortuosity. The graphs show the lack of significant difference in tortuosity (deviation the blood vessel length measured between two points on their curvature from the straight line drawn between these points) in the wild-type (blue dots and line) and nNOS KO (red dots and line) mice. n=12 analyzed fields from 3 mice for the wild-type and 10 fields from 3 mice for the nNOS KO. Data are presented in scatter plots (E) and relative frequency (F) **G** Analysis of the density of EdU-positive cells in the ependyma of the wild-type and nNOS KO animals. **H** Comparison of the fractions of the EdU-positive cells that are bound and not bound to the blood vessels (marked as BV-bound and BV-non-bound, respectively). n=42 analyzed fields of 150×150 micron collected from 3 mice for the wild-type and n=25 fields from 3 mice for the nNOS KO animals. I-N Density of B1 cells (I,J,M) and rosettes (K,L,N) in the ependyma of the wild-type and nNOS KO animals.

Furthermore, we determined the numbers of dividing cells in these brain regions by administering the thymidine analog 5-ethynyl-2’-deoxyuridine (EdU) 2 h before analysis. Thè total number of EdU-labeled newborn cells did not significantly differ between wild-type and mutant brains (Fig. 5G). We then analyzed subpopulations of EdU-labeled cells in the wild-type and nNOS KO mice according to their proximity to blood vessels, as the fractions of blood vessels-bound and blood vessels cells in the neurogenic area.

We found that in the wild-type group a substantial fraction of EdU+ cells ( 70%) were located near (1– 2 nucleus diameters) blood vessels, where they were born, while only 30% were found farther away from the blood vessels, being either migrated away from the blood vessels or born there by dividing neuronal progenitors.

In contrast, in the nNOS KO group the distribution between blood-vessels bound and blood vessels bound EdU+ cells was almost equal as increased fraction of EdU+ cells were located farther away from the blood vessels (Fig. 5H). This may be due to an alteration in the progenitor cell division, dynamics of lineage progression, migration, or in a combination of these processes. Together, these results indicate that nNOS has a role in the regulation of neurogenesis in the lateral walls of the ventricles.

The evaluated the density numbers of B1 stem cells and the rosettes surrounding them and again and did not observe significant differences (Fig. 5 I-N).

## DISCUSSION

The ependyma, a ciliary epithelium that lines the walls of the brain ventricles, has crucial roles in generating CSF flow and, consequently, nutrient distribution, waste removal, and overall homeostasis in the brain and central nervous system. The cilia are highly effective in mediating the CSF flow across the ventricles: in the mouse brain, only a few seconds are required for the CSF stream to traverse the lateral ventricle (8). This efficiency is supported by the concerted high-frequency beat of numerous cilia. It is also supported by the cellular architecture of the ventricular wall, where multiciliated cells are polarized and arranged to support the CSF flow. In particular, the translational polarity vectors, when analyzed for small groups of cells (12, 26), show alignment with the direction of the flow. However, the tissue-level polarity for large areas of ependyma, with hundreds and thousands of cells, as well as the correspondence between translational polarity and the flow remain unclear.

Our study, by integrating *in vitro* data of ciliated cell polarization across the ependymal tissue and live imaging of cilia motion, flow velocity, and trajectories in the lateral ventricles, suggests that the CSF flow, while maintaining a general P-A direction in the dorsal part of the ventricle, includes local microflows with both straight and curved trajectories that deviate from the main direction of the flow.

Our findings reveal that fluid flow in the lateral ventricles is more complex than previously understood, involving not just mainstream flow but also numerous microflows along the ventricular walls. These microflows likely arise from clusters of multiciliated cells with locally aligned translational polarity.

The demarcation between domains was evident when the flow of microbeads was examined, particularly in the areas where the microbeads’ trajectories abruptly changed direction (Figs. 4D-I, S1-S3). Interestingly, whereas most ciliated cells within a particular domain exhibited coordinated ciliary motion, we also observed isolated cells and groups of cells whose motion deviated from the predominant direction within their domain. However, these outlier cells did not appear to significantly interfere with the domain’s collective ability to produce a directed local flow. In addition, we identified groups of cells arranged in circles, with each cell’s ciliary motion oriented radially relative to the circle’s center (Fig.4E, S2). Analysis of the flow trajectories revealed that the path of the flow shifted direction when passing over these circular cell groups. We propose that these unaligned groups of cells, along with the defined domains, might function as “road signs” directing a “turnaround” or a “detour” flow path, thus contributing to the generation of complex microflows with both straight and curved trajectories. We further hypothesize that such microflows might be particularly important for sharply bending the otherwise linear flow of CSF, e.g., for directing the flow along the curvature of the anterior horn. Overall, the variety of ciliated cell arrangements generated a variety of local microflows along the lateral ventricular wall.

In parallel with analysis of flow dynamics in live preparations, we examined tissue-level translational polarity in fixed samples, assessing the variation of this feature *in situ* across extended ventricular areas. Our novel algorithm to evaluate translational polarity vectors provided insights into the vectors’ spatial distribution across the tissue, in contrast to the common statistical evaluation of this feature as circular plots.

Remarkably, by examining the distribution of the vectors in situ in these areas, we found that the alignment of translational polarity vectors occurred only among immediate neighboring cells within relatively small clusters of cells. For instance, an examined area sample 0.065 mm**^2^** was comprised of 14 such clusters, each with a distinct median translational polarity vector (Fig. 3K). This finding challenges the prevailing belief that translational polarity of ependyma is universally aligned along the mainstream flow.

Assuming that the translational polarity vector in a particular cluster is aligned with the local flow direction, we inferred that the cellular arrangement and cluster’s translational polarity supports the formation of multiple microflows in the ependyma with varied directions additionally to the major P-A CSF stream. This proposition was supported by the live imaging experiments and recordings of the composition of local microflows with varying directions and trajectories.

We also hypothesize that the arrangement discovered herein might have evolved to optimize the distribution of the mainstream of CSF across a wide surface area. We propose that local coordination of the ciliary motion in numerous domains of ciliated cells may have advantage over the uniform tissue-wide alignment of the ciliary motion by spreading the CSF further from the main stream, thereby ensuring thorough tissue cleansing and nutrient delivery. In addition, the presence of domains with local microflows may be important for bending the otherwise linear trajectory of the CSF flow.

Remarkably, we also found that ciliated cells, which are elongated at their apical surfaces, are aligned by their long axes of the apical surface and oriented almost orthogonally relative to the A-P axis of the brain. Whether this feature is associated with translational polarity and flow directionality is not yet clear.

Our conclusions are consistent with prior findings indicating a complex pattern of flow in the third ventricle, produced by interactions between straight and bent flows, supported by the variations in ciliary motion orientation (Faubel et al., 2016). Our results are also consistent with the observation that, in zebrafish nose epithelium, ciliary motion is coordinated only locally, without global tissue-wide coordination (Ringers et al., 2023).

The connection between local cilia architecture and the topology of the flow they generate remains poorly understood. Studies of mouse trachea found significant variations in cilia organization, with rotational polarity of the ciliated cells aligned only locally, with their immediate neighbors (28, 29). These studies also found numerous variations of the local flow patterns. Remarkably, despite these variations, the fluid transport remained coherent along the lungs–nose axis. A model based on these observations proposes how the variations-based noise can improve the overall function of the system (28, 29).

The mechanisms generating the complex arrangement of multiciliated cell domains remain to be established. An initial plan that determines the domains’ structure and arrangement and thereby the microflow trajectories might conceivably be established during embryogenesis, e.g., through the core PCP pathway factors and, possibly, their interactions with the A-P axis-determining growth factors. The final pattern might be potentially fine-tuned by the feedback of the mechanical pressure of the fluid flow on the structural polarization of the multiciliated cells (30)

Finally, our results reveal the importance of nNOS in controlling the CBF and planar polarity in general, as well as its special aspect - translational polarity, and also in controlling the flow dynamics, i.e., the velocity and trajectories of the local microflows. Remarkably, we found that CBF is associated with ciliary beat coordination. Specifically, we detected areas with severely diminished CBF in the nNOS KO ependyma and found that in such areas, a decrease in CBF was associated with loss of ciliary motion coordination within individual cells. This finding suggests that cilia must reach a threshold level of CBF to synchronize their motion and that nNOS/NO may be important for intracellular ciliary beat synchronization. The scale of the response to the deletion of nNOS was found to vary across the ventricular preparation from the same animal. In some areas of the nNOS KO mutant ependyma, we observed severe defects in planar polarity, cell size, and CBF, whereas in other areas we observed only marginal effects. The limited penetrance of the nNOS deletion effect within the same organism is not yet clear. NO produced by other forms of NOS, such as endothelial NOS of the blood vessels, which are unevenly spread across the tissue, might possibly compensate for the lack of nNOS; thus, the degree of compensation might be associated with the local architecture of the tissue or the path of the blood vessels. Compensation might also arise from increased activity of other nitrosylating enzymes or decreased activity of denitrosylases (Stomberski et al., 2019).

Among the limitations of the study, we should point to the sampling approach for analyzing the ventricular ependyma. An approach that would allow a comprehensive representation of the entire surface of the ventricular wall (e.g., employing light-sheet microscopy) may provide additional insights into the domain arrangement across the entire ventricle. Also, it would be advantageous to be able to determine the flow, translational polarity, domain organization, and CBF in parallel and *in vivo* (e.g., employing compound transgenic lines expressing markers of planar polarity).

## Materials and Methods

### Animals

Mouse maintenance followed the guidelines for the use and treatment of laboratory animals from the National Institutes of Health and all experimental procedures were approved by Animal Care and Use Committee of Stony Brook University. Wild-type and mutant animals were bred and housed at the animal facilities of SBU in a standard light-and temperature-controlled environment (12-hour light/dark cycle; light on at 7:00 AM; 21°C) with access to food and water ad libitum. Generation of nNOSKOex6 knockout mouse line (nNOS KO throughout the manuscript) representing a null mutation of nNOS gene was described previously (31). This knockout line, generated by deletion of the active site encoded by exon 6 of the mouse gene, lacks functional mRNA or protein, and unlike the widely used nNOS mutant line (Jackson Labs #2633), represents a true null mutant of the nNOS gene. In most experiments, the nNOS KO+/+ and nNOS KO-/- mice from the same litter were used as wild-type and knockout specimens. Brain specimens of nNOS-CreER::Ai9 transgenic mouse (25) after induction of recombination with tamoxifen was a gift from Dr. Josh Huang (CSHL).

### Dividing cell labeling

To label the dividing cells in the brain, animals received a single peritoneal injection of 5-ethynyl-2’-deoxyuridine (EdU) (Sigma-Aldrich), at 123 mg/kg 2h before the analysis. To reveal the labeled cells and the adjacent blood vessels, mice were perfused with TexasRed (Vector Laboratories) and the explants were stained through a click reaction in PBS containing 0.2% Triton X-100 (Sigma-Aldrich), 120 mM sodium ascorbate (Sigma-Aldrich), 6 mM CuSO4 (Sigma-Aldrich), and 10 µM AlexaFluor 555 Azide (Invitrogen) for 20 min at RT in the dark before stopping the click reaction by incubation with 0.1 M EDTA (Sigma-Aldrich) and washing in PBS as described (32–34).

### Immunocytochemistry and microscopy

We used following antibodies for immunocytochemistry: sheep polyclonal antibody to nNOS (ab6175 ABCAM), mouse monoclonal antibody to acetylated α-tubulin (Sigma T6793, dilution 1:1000); mouse monoclonal antibody to γ-tubulin (Sigma T6557, dilution 1:500); mouse monoclonal antibody to ZO1 (ThermoFisher 33-9100, dilution 1:200), rabbit polyclonal antibody to cenexin (ODF-2) (Sigma HPA001874, dilution 1:500). Secondary antibodies were anti-mouse, anti-rabbit, anti-sheep, or anti-goat antibodies conjugated to Alexa-488, Alexa-558, or Alexa-633 or Alexa 405 from Molecular Probes, used at dilution 1:500. Actin was visualized with phalloidin-Alexa 488 or phalloidin-Alexa 633 (Molecular Probes). Blood vessels were visualized with Dylight 649 or TexasRed (Molecular Probes). Lycopersicon Esculentum Lectin (Vector Laboratories, DL-1178) injected in tail vein or sub orbitally in mice at dilution 1:100.

### Whole mounts of the lateral ventricle wall

Craniotomy was performed to isolate the lateral ventricles as described (12). The resulting whole-mount preparations were then fixed in 4% paraformaldehyde in 1xPBS for 20 min followed by a wash with 0.1% Triton x100. Blocking of the samples took place for 1 h at RT or overnight at 4°C in a solution of 0.3% Triton-PBS and 10% BSA. The same blocking solution was used for both primary and secondary antibody incubation at 4°C overnight.

### Microscopy

Samples processed by IHC were visualized with confocal microscope Leica SP8X or SP5 II. Optic sections of 1024×1024 px were collected in steps 0.5-1 micron through the depth 2-5 micron, then reconstructed and presented as maximum projection images. Adjustment of brightness and contrast of the pictures, if required, was performed using Photoshop CS5 (Adobe Systems) or ImageJ software.

### Recording of cilia motion and flow

Freshly dissected and flattened brain ventricles were placed in DMEM high glucose solution, containing 1 µM DEEP Red 647 (Invitrogen T34077) and incubated for 1 h at 37°C. After that the specimens were placed into a custom-made chamber, where they were oriented stereotypically in regards to the A-P and D-V body axes. To record the flow generated by the cilia motion, fluorescent beads carrying Alexa 488 (2 μm FluoSpheres; Invitrogen) were added to the chamber. Live imaging was conducted with Zeiss Axiophot, retrofitted with a filters cube, allowing to segregation 488 and 647 emissions, and high-speed PRIME LED camera. Movies were recorded at x20 and at x40 magnification for 3-4 sec at 65-120 fps with ImageJ and Micromanager software.

### Statistical methods

Biotool1 (github.com/pol51/biotool1) was used to analyze polarity changes at the tissue- and cellular level(26). Tissue-level changes were measured using the angle representing the radial difference between the mean vector of the field and the vector of the patch of an individual cell. Resulting angles of tissue level displacement obtained from biotool1 software were transformed into circular vectors using Oriana software. The patch angle indicates the intracellular coordination of patch displacement. The expected pattern of an angle for a wildtype ciliated specimen shows range within 45° clockwise or counterclockwise (26). Cell-level changes are represented by a patch displacement value which is calculated as normalized quotient/ratio of patch distance between the cell center and center of patch to the distance from the cell center to the membrane. A smaller quotient corresponds to a patch closer to the perimeter of the cell, indicating a greater asymmetry. The length and tortuosity of blood vessels was determined using “Skeletonize (2D/3D)” plug-in for topological skeletonization in ImageJ (https://imagej.net/Skeletonize3D).

The statistical ANOVA test, t-test, and Wilcoxon test performed in Excel, Image J, Oriana, and GraphPrism. Graphs for the analysis were made in GraphPrism and Adobe Illustrator. Graphs are presented as mean+/- standard error of the mean. Statistically significant differences are indicated (*, P < 0.05; **, P < 0.01; ***, P < 0.001).

### *in situ* CBF determination

We developed a program that estimates the pixel-wise cilia beating frequency (CBF) in videos of ciliated cells. Before we measure CBF, we needed to preprocess the videos to remove undesired noises which can disturb the measurement of the true CBF in videos. By observation, the noise in the videos is salt-and-pepper noise. Therefore, we applied a median filter to video frames to remove the noise. Specifically, the median filter runs through each frame pixel by pixel, replacing each pixel with the median of neighboring pixels. Given a denoised video of size T×H×W, we computed the Discrete Fourier Transform of every spatial pixel. We regard the frequency with the maximal amplitude as the dominant beating frequency in each pixel. After we obtained the pixel-wise beating frequency map, we needed to post-process the map to remove out-of-the-focus area predictions and non-ciliary area predictions. We hypothesize that the beating frequency in these areas has a smaller amplitude. Thus, we set a threshold to filter out the areas where the beating frequency amplitude is smaller than N. The value of CBF in each cell was represented in the heat map as a fraction of 100%. The algorithm is in the link to be posted in the supplement.

### Orientation of the ependymal ciliated cells relatively to the A-P axis

We approximate each cell as a sequence of points along the cell boundaries. The longest axis is the longest distance of a pair of boundary points. For computational efficiency, we computed the convex hull (a polygon) that includes the cell tightly. Then, we looped over the convex hull vertex pairs to identify the longest vertex pair distance. After the longest vertex pair was located, the longest axis was the segment that connects these two vertices. The algorithm is in the link to be found in the supplement.

### Translational Polarity

The vectors of translational polarity were calculated based on the images sampled across the dorsal area of the ventricular wall of each animal in groups of at least 5 littermates with wild-type or nNOS genotype. For each cell and the patch within the cell, we computed their geometry center respectively. Then, the vector pointing from the cell center to the patch center represents the translational polarity direction. The algorithm is in the link to be found in the supplement.

### Tracking beads movements

The beads in the first frame are denoted with red color while the beads in the last frame are denoted with green. For the intermediate frame, we used a mixed color which gradually transitions from red to green. Then, we averaged the beads of all frames which produced a track for each bead. In the figures, the bead track direction is indicated by the red-to-green transition direction.

### Cell clustering by translational polarity vector

We used K-means to cluster the cells into domains in each of which cells manifest similar translational polarity directions. Specifically, K-means algorithm is an iterative clustering technique that partitions a set of data points, i.e., cells in our case, into K clusters by minimizing the sum of squared distances between data point features and their corresponding cluster centroids. To increase the spatial continuity within each domain, we build the feature of each cell by combining its translational polarity direction and its spatial location. Then, we perform K-means repeatedly with different number of clusters. We select the number of clusters for final visualization by thresholding the resulted standard deviation of translational polarity angles.

## Supporting information

Supplemental movie S4 WT cilia motion

Supplemental Movie S7 nNOS KO cilia motion

Supplemental movie S8 WT cilia motion High Mag

Supplemental movie S9 nNOS KO cilia motion

Supplemental data description

## Acknowledgments

We thank Vladimir Maxakov and Evgeny Amelchenko for help with the experiments. This work was supported by grants from the National Institute of Aging: R01AG AG057705 to G.E., H.B., and N.P., R01 AG076937 to G.E. and N.P., and R01 AG076937-S1 to K.M.; grant from National Heart, Lung, and Blood Institute (5P20HL113443-02) to G.E. and N.P.; SBU SEED grant to N.P and D.S.; and intramural departmental grants to N.P.

## Supplementary materials

### Supplementary figures

**Supplementary Figure1/SF1**

Expression of nNOS in the nNOS-CreER/Ai9 mice. dTomato expression, driven by nNOS promoter, is observed in the ciliated cells, with the sample counterstained with antibodies to anti-dTomato, acetylated α-tubulin (red) and actin (blue). Scale bar is 10 µm for (A-C). After inducing nNOS-driven recombination with tamoxifen, dTomato signal is observed in the ependymal cells of the lateral ventricular wall. Co-staining with antibodies to dTomato, acetylated alpha-tubulin (a ciliary marker) and phalloidin (to highlight the apical actin cytoskeleton) revealed dTomato signal on the apical cell surface of the ciliated cells, concentrated in the tufts of cilia, implying nNOS expression in the multiciliated cells of the ventricles

**Supplementary Figure 2 /SF-2**

nNOS deletion does not cause defects in the basal bodies docking

## Supplementary movies

S1-S3, represented as frames in Figure 4, cilia motion in wild-type S4 Cilia motion in the ependyma of WT

S5-S6 represented as frames in Figure 4, cilia motion in nNOS KO S7 Cilia motion in the ependyma of nNOS KO

S8 Cilia motion in WT, high magnification

S9 Cilia motion in nNOS KO, high magnification

