## Supplementary figures and images for "Ciliated cell domains with locally coordinated ciliary motion generate a mosaic of microflows in the brain’s lateral ventricles"

### Supplemental Figure1

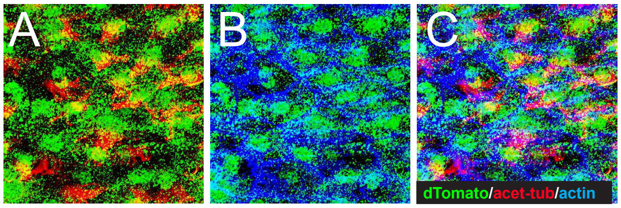


Supplementary Figure 1

### Supplemental Figure 2

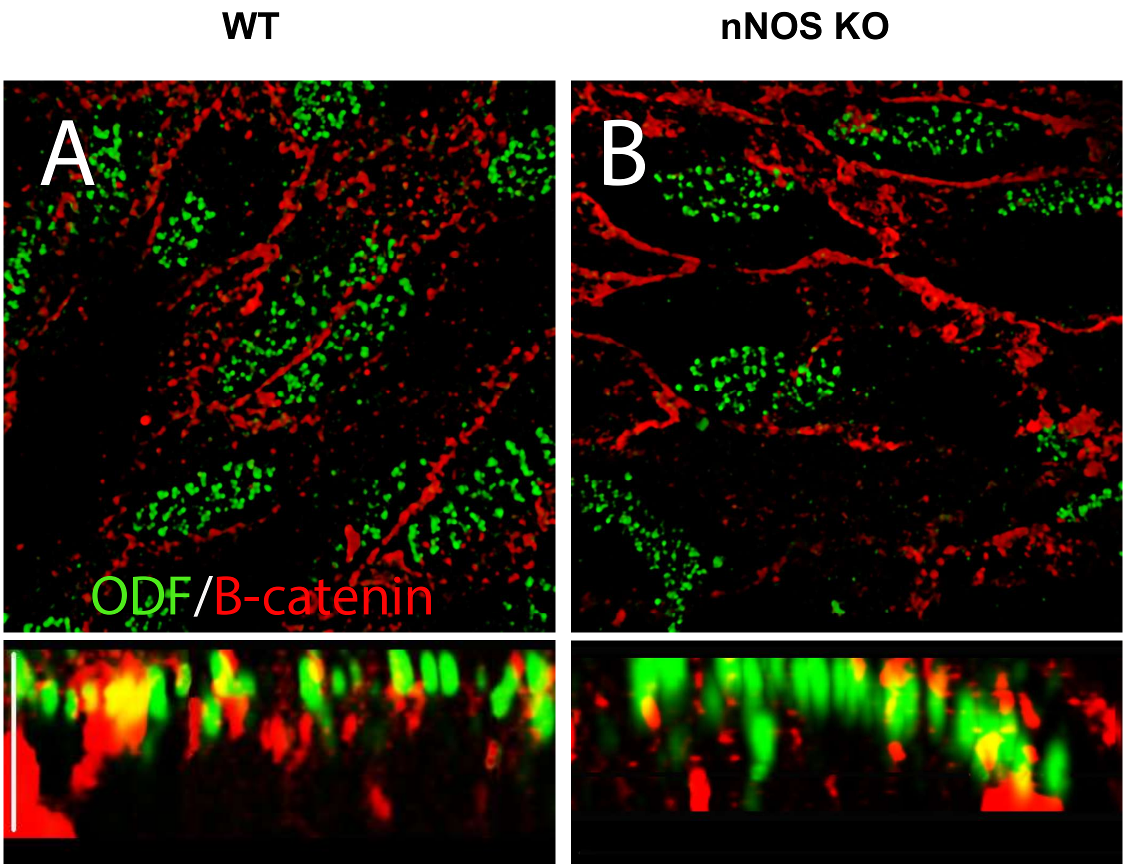


Supplementary Figure 2
